# What can an evolutionary perspective on women’s health provide to the current public health paradigm? A case study on redefining thyroid cancer as a reproductive cancer

**DOI:** 10.1101/222737

**Authors:** Abby Fraser

## Abstract

Incidence rates of non-communicable diseases in women are increasing worldwide and now contributing more to mortality and morbidity than reproductive health. Despite this universal trend, there is no recognition by the WHO of this phenomenon in women’s health. Additionally, thyroid cancer is considered a non-sex specific cancer despite female incidence rates being triple those of males. To understand why there are such conceptual limitations to women’s health, biomedicine - and its prevalence in western public health authorities - will be analysed from an anthropological perspective. In mind of these conceptual limitations, insights from evolutionary perspectives on health are introduced as well as how they may alleviate the conceptual barriers in the current biomedical paradigm. Using the example of breast cancer, the difference between a reproductive cancer in women, and cancer located in the reproductive tract will be explored. From this, the possibility that a cancer outside the reproductive tract can be directly influenced by reproductive function is explored in the specific case of thyroid cancer. Thyroid cancer and current public health approaches to preventing malignancies of the thyroid are examined to show the limitations of their scope. Furthermore, evidence supporting the direct link between thyroid function and reproductive function is presented. Current academic studies into the link between thyroid cancer and women’s reproductive function are analysed to show they are subject to the same conceptual limitations of women’s reproductive function as found previously in biomedicine. To conclude, I will introduce a new hypothesis for exploring the impact of women’s reproductive function on thyroid cancer incidence rates. This hypothesis will allow women’s health to be viewed holistically, and allows reproductive function to be investigated beyond parity. Thus, the conceptual limitations of women’s health in the public health paradigm will be alleviated.

## 1. Introduction

The United Nations (UN), World Health Organisation (WHO) and United Nations Population Fund all approach women’s health problems from the same perspective. The 2009 WHO report on ‘Women and Health’ directly states that

“…women and girls have particular health needs…that only women experience and that have negative health impacts that only women suffer”. (P.3)

These ‘particular health needs’ are those stemming from women’s reproductive function. Access to contraception, adequate antenatal care and appropriate parturition care are considered aspects of women’s health and their importance to intergovernmental health policies is cemented in their inclusion in the Millennium and Sustainable Development Goals. These are goals established by the UN in 2000 and 2016 respectively as aspirational targets in the campaign against eradicating poverty (Table 1.1).

**Table 1.1.**
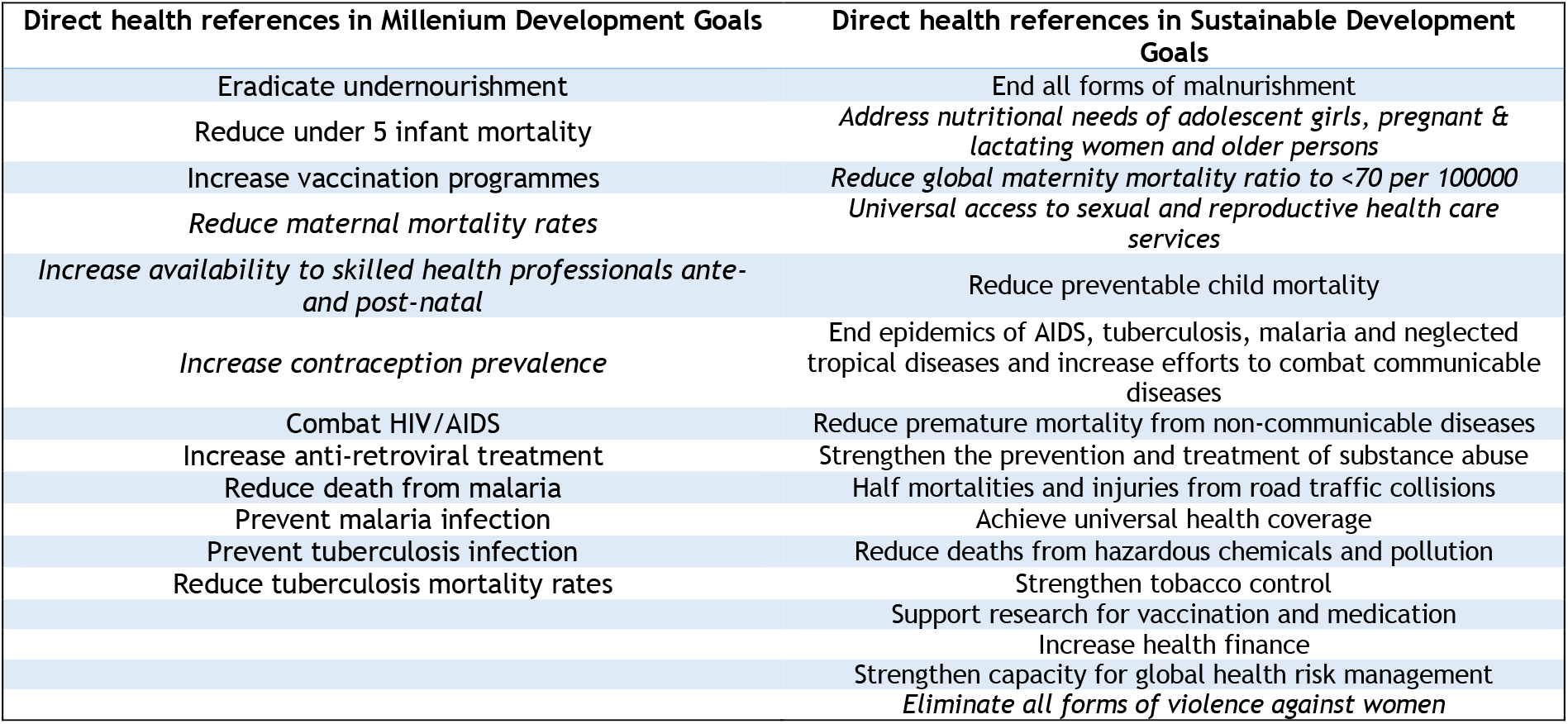
All UN development goals directly related to health. Goals directly related to women’s health have been italicised (UN 2015, UN Women 2016)

| Direct health references in Millenium Development Goals | Direct health references in Sustainable Development Goals |
| --- | --- |
| Eradicate undernourishment | End all forms of malnurishment |
| Reduce under 5 infant mortality | *Address nutritional needs of adolescent girls, pregnant & lactating women and older persons* |
| Increase vaccination programmes | *Reduce global maternity mortality ratio to <70 per 100000* |
| *Reduce maternal mortality rates* | *Universal access to sexual and reproductive health care services* |
| *Increase availability to skilled health professionals ante- and post-natal* | Reduce preventable child mortality |
| *Increase contraception prevalence* | End epidemics of AIDS, tuberculosis, malaria and neglected tropical diseases and increase efforts to combat communicable diseases |
| Combat HIV/AIDS | Reduce premature mortality from non-communicable diseases |
| Increase anti-retroviral treatment | Strengthen the prevention and treatment of substance abuse |
| Reduce death from malaria | Half mortalities and injuries from road traffic collisions |
| Prevent malaria infection | Achieve universal health coverage |
| Prevent tuberculosis infection | Reduce deaths from hazardous chemicals and pollution |
| Reduce tuberculosis mortality rates | Strengthen tobacco control |
|  | Support research for vaccination and medication |
|  | Increase health finance |
|  | Strengthen capacity for global health risk management |
|  | *Eliminate all forms of violence against women* |

The goals pertaining to women’s health specifically in the Millennium Development Goals focused on reducing maternal mortality rates and improving access to family planning methods and care throughout pregnancy. Efforts to improve women’s health needs further were included in the Sustainable Development Goals with the continued goal of reducing maternal mortality and increasing access to family planning advice and methods. However, there has been no recognition of non-communicable diseases falling under women’s health needs.

The disparity between the perception of ‘women’s health needs’ and actual women’s health concerns is beginning to be criticised. In 1990, 4 out of the 10 leading causes of women’s death were related to reproductive and maternal mortality but in 2013 7 out of 10 leading causes of women’s death worldwide were caused by non-communicable diseases (Peters, Woodward et al. 2016). The 2016 ‘Women’s Health Agenda’ was published by the Oxford Martin School as a policy recommendation focusing on the impact of non-communicable diseases to women’s health and wellbeing globally. The Agenda criticises the UN and WHO perspective, especially the consistent focus on improving sexual and reproductive health (Peters, Woodward et al. 2016).

Thyroid cancer is the most common endocrine malignancy worldwide and the fifth most common cancer in women (Nielsen, White et al. 2016, Zahid, Goldner et al. 2013, Cabanillas, Mcfadden et al. 2016). While considered a non-sex specific cancer, incidence rates in women are three times higher than those in men, with incidence in both increasing overall. This follows the general trend in the rise of non-communicable diseases as prominent causes of morbidity and mortality for men and women worldwide. Despite the growing prominence of non-communicable diseases as women’s health problems worldwide, there is little or no addressing of this phenomenon in the universal public health approach to women’s health. To explain this, I will first explore why women’s health is not able to encompass non-communicable diseases. I will then introduce a framework to health that allows non-communicable diseases, including thyroid cancer, to become an issue of women’s health.

First, I will begin by analysing why these conceptual limitations of women’s health exist within medicine and public health as disciplines. Biomedicine and the biomedical perception of health will be investigated to identify the main conceptual limitations to women’s health in public health institutions. The creation and perpetuation of these limitations in Western public health institutes will be discussed. As biomedical frameworks for approaching women’s health and health overall contain conceptual limitations, there is a strong case for introducing a new framework for health.

Secondly, I will provide justification for the application of evolutionary theory to health and disease as a mode through which these conceptual limitations can be alleviated. Evolutionary theory, an emerging discipline, conceptualises health as an optimum state for reproduction. Using the example of breast cancer, malignancies of the reproductive system will be shown as not always reproductive in nature. The framework of women’s health and disease is then expanded to consider the influence of reproductive function on non-reproductive malignancies.

Thirdly, I will focus on thyroid cancer, presenting its current epidemiological trends and how it is approached by public health authorities. I will then present evidence supporting an inherent interconnection between thyroid function and women’s reproductive function. From this I will criticise current academic approaches to understanding the link between women’s reproduction and thyroid cancer incidence rates. From here, I will propose a new hypothesis linking increased risk of thyroid cancer with a woman’s reproductive function. Through this hypothesis thyroid cancer can be redefined as a reproductive cancer.

By approaching the current biomedical discourse in public health from an anthropological perspective, I will identify the conceptual limitations to women’s health present in dominant public health institutes. I will then introduce an evolutionary approach to health as a solution for alleviating these limitations. I will conclude with the case study of thyroid cancer, showing how conceptual limitations emerge practice within current public health approaches to the disease. Using insight gained through an interdisciplinary approach to women’s health, I propose a hypothesis which encourages a non-discriminatory view of women’s health and its needs.

## 2. What are the conceptual limitations to women’s health in public health discourse?

### Synopsis

Drawing on current literature from medical anthropology and related fields, I have identified the three main limitations of public health approaches to women’s health - the reduction of women’s health to reproductive function; the inherent pathologisation of women’s health; and the biomedical hegemony within public health institutions which perpetuates limited research scope as well as the systematic exclusion of women from clinical research. Reduction of women’s health to reproductive function occurs through its entwinement with demography, causing women’s health to encroach on demographic, social and political dimensions. The inherent pathologisation of women’s health is directly related to the biomedical tenet of the soma, which relates to how health and disease are conceptualised in the biomedical paradigm. Finally, the hegemony of biomedical institutes in public health will be explained in relation to the biomedical tenets of causation in health and disease. The prominence of biomedical frameworks in public health implicitly cause limitations to women’s health research and research into non-communicable diseases on a whole.

### 1.1 Reproductive essentialisation of women’s health

The first limitation to women’s health research exists in a reductionist view whereby women’s health is equated to healthy reproductive function. The modes through which this is achieved will be discussed. As mentioned, the specific women’s health needs identified by the WHO involve those pertaining to women’s reproductive function. This initial reproductive essentialisation of women’s health, i.e. the generalised focus on the women’s reproductive function as an indicator of health, results in their health becoming of demographic interest. The current WHO definition of reproductive health states it to be:

“A state of complete physical, mental and social well-being and not merely the absence of disease or infirmity, in all matters relating to the reproductive system and its functions” (UN 1995): p.59)

‘Matters relating to the reproductive system’ is expanded to cover reproductive factors such as a satisfying and safe sex life; the capability to reproduce and the freedom to decide if and when to do so; the rights to be informed of family planning choices; and the right of access to appropriate healthcare services during pregnancy and childbirth (UN 1995). Three dimensions of reproductive health therefore exist - the human condition related to individual health and wellbeing; an approach for policies, legislation and attitudes; and as services, their utilisation and their provision (Ritu 2002). In doing so, a woman’s reproductive function becomes inextricably linked with the formation of family planning policies and the services it requires. This extension conflates the women’s reproductive tract to all social, healthcare and population concerns within the society in which the woman exists. Furthermore, the dualistic view of the woman as either being in a state of fertility or infertility, either due to family planning methods or pregnancy, reinforces the role of the woman as a reproducer in a medical context. The medical definition and scope of reproductive health is therefore as such - morbidity and mortality related to giving birth and being a mother. This reductionist view of women’s reproduction and the reproductive tract allows it to be considered a separate reproductive entity to all other physiological mechanisms within the women’s body and that all causes of morbidity in the reproductive tract come from the processes of sex and childbirth. Other conditions afflicting parts of the body outside the reproductive tract (aside from the breasts) will not have a condition of the reproductive tract favoured in their aetiology (Qadeer 1998).

The WHO definition of reproductive health includes the statement that reproductive health encompasses the capacity to reproduce and the freedom to decide if, when, and how often to do so (UN 1995). By bringing in the concept of fertility and the number of children a woman may have, reproductive health begins to encroach on multiple disciplinary boundaries such as those of demography, human rights and social development (Wang 2016). The contemporary issue of population growth has subsequently become one of fertility control in many regions around the world and, due to reproductive essentialism, women’s health becomes embroiled in policies designed to control population (Wang 2016). Combined with the multidisciplinary recognition of the positive effect of family planning on maternal and child health, the importance of women’s access to contraception on controlling fertility levels has caused family planning to be viewed as a vital preventative and positive health measure (Wang 2016, Dixon-Mueller 1993). This demographic perspective of reproductive health is grounded in the neo-Malthusian thought in which contraceptive effectiveness and preventative checks are incentivised for reducing population growth (Wang 2016). Given that contraceptive availability is included in the WHO definition for reproductive health, demography has intertwined itself in to the agenda for women’s reproductive health. The relationship between population and women’s reproductive health also extends into political realms given that many states’ power has depended directly and indirectly on defining the normative family of the nation-state and controlling populations (Ginsburg, Rapp 1991). This is enforced by the recognition of population change, labour markets and economic growth of a country being connected and influenced directly by population control (Ginsburg, Rapp 1991). Thus, social movements concerning various aspects of women’s health and reproduction can occur in many contexts including social justice, population control and economic change (Ginsburg, Rapp 1991). The perceived importance of the women’s reproductive tract therefore becomes intrinsically linked to social movements striving towards the goal of modernisation and societal change.

The reproductive essentialisation of women’s health is deeply engrained within public health discourse to the point that healthcare provisions for women of reproductive age in many populations are solely concerned with women as reproducers and as mothers to children (Inhorn 2006). Treating reproductive health as a discrete entity within the individual woman results in reproductive essentialisation in which multiple dimensions of women’s health are neglected in clinical and research contexts (Inhorn 2006, Qadeer 1998). In addition to the separation of reproductive function from general health, by defining a woman to be of reproductive age compartmentalises women’s lives and health and creates artificial boundaries within the continuity of a woman’s life (Qadeer 1998). Thus, deeper insights into the health of women and connections between different physiological properties are not discovered because of their low priority in a research setting. Essentialisation causes a measurement trap in which a lack of information from research about women’s health on a whole is both the cause and effect of continuing ignorance of the subject (Graham 1998, Graham, Campbell 1992). As long as women’s health is reduced to parturition and procreation, there is little requirement for the effect of reproductive function on non-reproductive processes in the body to be investigated. In a self-fulfilling prophecy, public health will continue to consider reproductive health to be a primary concern for women around the world due to the lack of evidence for other perspectives into health and wellbeing for women. Criticisms of this reductionist view, such as the Women’s Health Agenda (Peters, Woodward et al. 2016) are beginning to emerge and are highlighting this skewed perception of women’s health. To identify other areas of health and wellbeing in women’s which can be approached from population level policies, a conceptual shift towards viewing women’s health in its entirety and outside the context of pregnancy must be made in women’s reproductive health and women’s health overall.

### 1.2 The inherent pathologisation of women’s health

A major limitation of the biomedical frameworks in public health research specifically involving the women’s body is the systematic exclusion of females from all levels of biomedical research. Given the importance of positivistic, systematic objectivity in biomedical research, which will be explained shortly, the pathologisation of women’s reproduction is often used as a valid reason for the exclusion of females from biomedical studies ranging from cellular experiments to clinical trials. The unpredictability of women’s health as perceived by the biomedical model justifies its exclusion on the grounds of ‘ensuring controllability’ within test subjects (Mazure, Jones 2015). By perpetuating the idea that women’s health is inherently unhealthy and prone to dysfunction, the use of all-male test subjects becomes an appropriate solution to controlling for health in the sample. The systematic exclusion of females occurs throughout preliminary cellular and animal trials in addition to human trials and thus, the assumption that the women’s body is a deviation from the male soma persists within biomedical research (Johnson 2014).

To explain this phenomenon in biomedical research, the inherent pathologisation of women’s health must be explored. This stems from the fundamentals of the biomedical approach to health, particularly the ‘soma’ which will be introduced subsequently. As mentioned, women’s health needs from a public health perspective are those which “only women experience and that have negative health impacts that only women suffer” (World 2009b). In the context of public and global health policy formation matters of women’s health and women’s health needs are specified as those directly related to reproductive health and its function/dysfunction. More importantly, women’s health needs are related to dysfunctions of the reproductive system. High maternal mortality rates are used to justify increasing in the presence of skilled health professionals before and during birth allows. Maternal mortality is arguably the largest dysfunction of the reproductive tract - one that causes death - and can result from complications during childbirth either due to the mother’s health or the environment in which parturition is taking place. Even within family planning advice and contraceptive availability, advice is given pertaining to the health consequences of contraceptive use and controlled fertility for women. Only two perspectives are considered - the health consequences of not being pregnant (ie. the use of contraception and pregnancy prevention) versus the health consequences of being pregnant (Dixon-Mueller 1993). Judgement on whether to use contraceptive methods is thus based on the risks to health that either pregnancy or preventing pregnancy incurs. The pattern of women’s health from the public health perspective indicated that women’s health is both specific to the reproductive tract and inherently pathologised. Development goals concentrate on improving the health of the woman rather than maintaining her wellbeing, and only when she is of reproductive age. To investigate the origins of these limitations in the perception of women’s health in public health discourse, the role of the biomedical perspective in the inherent pathologisation of women’s health will be explored.

The central tenets of biomedical theory - atomism and the soma^1^ - are central to the inherent pathologisation of women’s health. Atomism is fundamental to the biomedical view of health, and postulates that nature, society and the body are separate discrete entities of life and that all systems within each can be reduced to the pure level of natural law (Davis-Floyd 2003). Within the ‘atom’ of the body, subsequent reductionism occurs to distinguish separate factors of the body and health from one another. Removal of rationality, culture and anatomical differences between humans allows the pure physiological and biochemical processes of the body can be objectively studied (Gordon 1988). This reinforces the idea that all organs and bodily systems are autonomous units within the person. To separate and reduce all functions of the body in this way allows for biomedicine to consider itself separating the ‘facts’ of the body from ‘values’ imposed by non-biological confounders (Gordon 1988). To achieve neutrality and universality in laws of health and disease is to achieve the positivistic view of health - that biochemical and physiological mechanisms will follow a law in response to a stimulus. Thus, medicine and the act of treating illness becomes a universal constant for physicians, allowing one treatment to be applicable for any individual. Within the specific context of the women’s body, this allows for the reproductive tract to be envisioned as an autonomous unit of the women’s anatomy.

In addition, the importance of discrete, quantifiable entities in biomedicine plays an important role in the definitions of health and disease. Health is based on a disease/non disease model and preferred boundaries between the two are those which are discrete, objective, valid and reliable (Kaufert 1988). The biochemical and physiological mechanisms that underlie the physical manifestations of disease in the body can be identified as pathological, with each biomedical process being considered a ‘law’ of disease. The ontological representation of disease - being able to identify a pathological entity is to be able to foresee a medical action - reinforces the positivistic goals of biomedicine and equates identification of the cause of disease to an effective cure (Canguilhem, Fawcett et al. 1978). To restore health these pathological processes should be eradicated (Wang 2016). From the disease/non-disease model promoted by atomism, health is considered as the condition of being that remains after illness and dysfunctions are factored out of the individual (Wang 2016).

The cumulation of these principles results in the creation of the biomedical soma-the universal somatic body in which health and disease are found as opposite poles along a biological continuum. It is in this principle that the inherent pathologisation of women’s health occurs. The laws of biomedicine which cause manifestations of the disease are found on the negative end of the biological continuum and through their treatment the soma can be brought back to the neutrality of health. Canguilhem specifies the importance of this perspective in asserting the power of biomedicine as a force to restore health to the individual. Disease is not simply a disequilibrium of the soma but a disequilibrium enacted by nature upon the physiological function of the body, through factors such as pathogens or environment, to instil a new unhealthy equilibrium in man (1978). From this perspective, nature cannot be trusted to maintain the healthy soma and the application of biomedicine is the only way in which the perfect soma can be maintained (Canguilhem, Fawcett et al. 1978).

Health conceptualised as a deviation from the norm is especially relevant to the perspective of women’s reproductive systems in the biomedical perspective. The importance of deviation in restoring health places pathological phenomena as nothing more than quantitative variations in physiological functioning (Canguilhem, Fawcett et al. 1978). From this perspective, there is no variation by nature which does not constitute a degree of pathology highlighting the importance of perfection in the soma. The ideal soma of health is based on the male body, causing variations from the ‘perfect’ male soma to constitute pathological variations. These variations are epitomised in the women’s reproductive anatomy and function, as identified through ethnographic research on pregnant women in the hospital environment, (Davis-Floyd 2003). In a clinical setting the extreme anatomical deviations of the women’s body from the male and the uniquely women’s biological processes of menstruation, pregnancy and menopause are presumed to inherently ‘malfunction’ (Davis-Floyd 2003). This strongly suggests that the pathologisation of women’s health in a public health context, though implicit, stems from biomedicine and its social construction. The dogmas of biomedicine also contribute to the maintenance of women’s health as a pathological variation throughout the history of the discipline in that pathological fact accepts explanation more that it stimulates questioning (Canguilhem, Fawcett et al. 1978). Disease becomes a proxy through which the proper physiological function can be revealed. Thus, pathologies are not questioned as it is health that is learned through their presence. By accepting that women’s reproduction is a pathology causes the persistence of this perspective in the biomedical paradigm and all disciplines that extend from it. Thus, to remove this pathologisation, the fundamental truths of biomedicine and their place in the discipline of public health must scrutinised.

### 1.3 Biomedicine as an incomplete framework

The conceptual limitations previously explained become pervasive in biomedical public health discourse due to the social construction of its being. Biomedicine is a dominant perspective in health and disease and thus its prominence in research perpetuates the limitations it confers. To understand how biomedicine has achieved its place of prestige in public health authorities, the biomedical tenet of causation must be addressed. The importance of causation follows from a positivistic approach to human health. Biomedicine as a discipline originated from the identification of pathogenic sources of disease (Lock, Nguyen 2010). This cemented the central tenet that specific disease states are achieved only by necessary causal factors (Lock, Nguyen 2010). Disease aetiology in biomedicine emphasises the importance of causal factors affecting biochemical and physiological processes in the body, whether these be pathogenic, environmental or sociological (Lock, Nguyen 2010). The success of this approach, relative to other medical approaches, in identifying the cause of infectious disease and subsequent eradication campaigns has also granted the rational, systematic objectivity in biomedicine therapeutic efficacy over a broad range of other medical systems (Lock, Nguyen 2010). Combined with the perfectionism of the soma, eradicating the cause of deviant pathologies on a population level allows health to be achieved.

Public health, based on the biomedical conceptualisation of health, becomes primarily associated with the eradication of deviant pathologies. This works to achieve the goal of reducing disease incidence in a population but it does not compare fully to the definition of health given by the WHO. By their definition, health is:

“a state of complete physical, mental, and social well-being and not merely the absence of disease or infirmity” (World 2009a)

With biomedicine associated with attaining a physical state of ‘perfection’, there is a discordance between the two definitions. The priorities of the WHO, responsible for monitoring and improving health worldwide, are both the eradication of disease as well as the maintenance of well-being. However, biomedicine, the predominant discourse in research used to inform healthcare policies, is solely concerned with the achievement of health. Health cannot be achieved in the biomedical model in the presence of negative health outcomes - death, disease, disability, discomfort and dissatisfaction (Graham 1998). On the positive end of the health outcome spectrum lie satisfaction and fulfilment - two abstract concepts of wellbeing that are not explicitly recognised (Graham 1998). Wellbeing as a human state is neither a discrete entity nor is it associated with a compartmentalised physiological function and thus cannot be an interest of biomedicine. Furthermore, wellbeing accepts there can be variations in the healthy body which opposes the biomedical notion that perfection as health does not accommodate variation. These positive indicators of health are required by the WHO to class an individual as ‘healthy’ but this is not required in the biomedical paradigm. As many epidemiological investigations into health focus on negative biological outcomes - the product of biomedical laws - as a quantitative source of data, there is a fundamental limitation in the biomedical perspective of health directly affecting its assessment from a public health perspective.

Aside from the conceptual limitations of the biomedical perspective, functional limitation can be found in the dominance of biomedical research in funding circles and public health academia (Gabbay 1999). This perpetuates the limitation of biomedical research regarding women’s health and non-communicable diseases. As mentioned, the importance of causation in biomedicine stems from its initial application to treating pathogenic diseases. While risk of acquiring infection is largely determined by sociocultural factors in the individual’s environment, the causative factor of infectious diseases is pathogen introduction to the body. However, the dominant cause of mortality and morbidity in many societies comes from non-communicable or chronic diseases. Degenerative and non-communicable diseases differ from infectious diseases on one main point - the conditions of their development in the body are multifactorial. Unless a disease has a high genetic heritability, the risk factors for its development will be numerous and most likely associated with the socioeconomic environment in which the individual lives (Gabbay 1999). It is in the inherent multifactorial nature of non-communicable diseases that biomedical discourse severely limits the research performed by public health academics. As fewer and fewer biomedical causes for non-communicable diseases are found, given the strong biocultural component of these conditions, the current biomedical paradigm exhausts research elucidating molecular and cellular mechanisms underpinning non-communicable diseases (Gabbay 1999).

Bias towards biomedical research impacts the establishment of interdisciplinary biocultural research studies to be made, reducing studies focusing on novel approaches to public health questions (Gabbay 1999). Thus, despite efforts to expand public health research to address women’s health needs and particularly the impact of non-communicable diseases on women’s health, biomedical institutions will not enable such research to be funded or conducted. This is attributed to the co-evolution of public health as a discipline with biomedical studies, causing students and academics to naturally follow well established biomedical paths in research. Interdisciplinary studies lacking the establishment and reputation of biomedicine are not easily facilitated by public health institutions. Within the UK, most funding for public health programmes comes from NHS research and development programmes, the medical research council and medical charities (Gabbay 1999). The hegemony of biomedicine in the UK is also largely caused by the formation of the NHS. Politicisation of healthcare as caused by the welfare state creates structural barriers to interdisciplinary research. Power over of health policy has been ceded by politicians to the medical profession given their importance in the running of the NHS (Quirke, Gaudillière 2008). This places professionals with a biomedical background in positions of authority regarding public health funding (Quirke, Gaudillière 2008). Empowerment of clinical medics and academics with biomedical backgrounds has allowed the hierarchy or biomedicine over biocultural approaches to be maintained. As long as this hegemony persists, structural barriers will exist for conducting interdisciplinary research - especially that pertaining to women’s health.

### 1.4 Assessment of limitations: Do current critics provide adequate alternative frameworks?

Despite discussing the multiple limitations present in modern biomedical discourse, this does not mean that public health authorities have exhausted their means of researching and informing policies. Other conceptual approaches to health and disease offer different frameworks in which research surrounding non-communicable diseases and their risk factors can be studied. In this section, I will focus on two frameworks which may alleviate the conceptual and tangible limitations to research found in biomedicine.

One such framework is the ecological approach to health and disease as outlined by McElroy and Townsend (2004) which conflates health to one measure of environmental adaptation (Figure 2.1). The environment is formed of the biotic (e.g. food, pathogens, predators), abiotic (e.g. climate, materials, energy) and the cultural (e.g. socioeconomic position, technology, ideology). The interplay between culture and human biology, and thus the human’s ability to adapt, is crucial for understanding the health of both the individual and the population. Imbalance of any variable in the ecological approach to health is associated with disease and stress either directly or indirectly (McElroy, Townsend 2015). Rather than disease being viewed as a deviation from the norm within the body, it can be better understood as the result of environmental deviations or imbalances. Given the importance of socioeconomic factors to the aetiology of many non-communicable disease, a biocultural model which incorporates exogenous factors such as food availability, socioeconomic structure and physical environment is better suited to exploring incidence rates from a public health perspective. While this approach does address the need to apply a holistic perspective on health and public health policy it does not address the inherent pathologizing of women’s health or its reduction to reproductive function.

**Figure 2.1:**
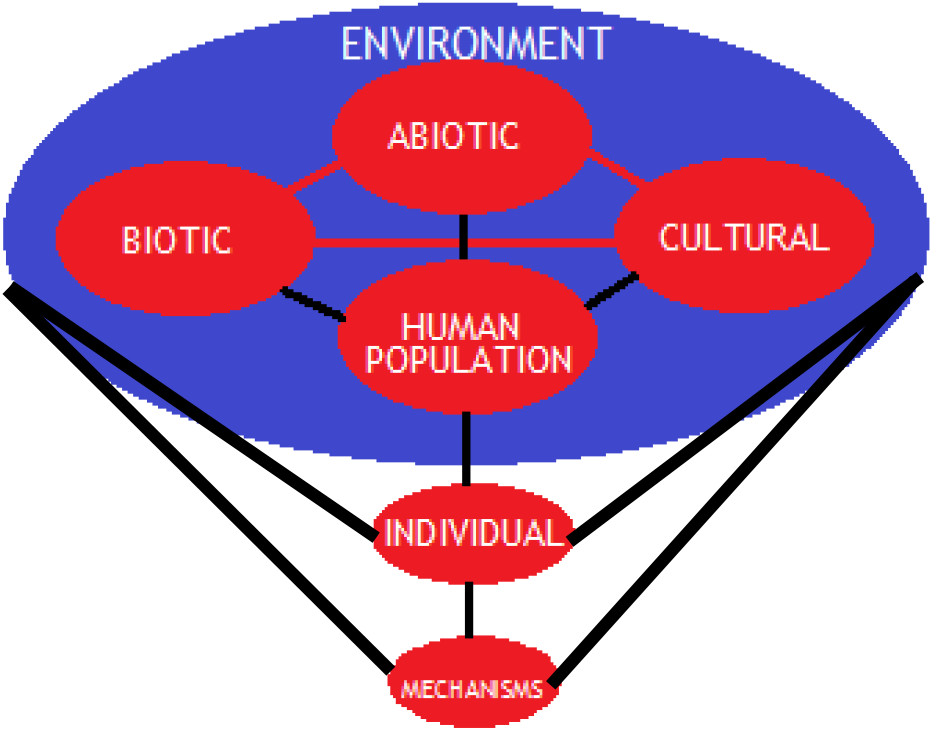
Human ecological model showing the interactions between the individual and the environment. Adapted from McElroy & Townsend (2004)

Expansion of women’s health to encompass both reproductive and non-reproductive health needs is proposed in the Women’s Health Agenda (2016). The agenda, which recognises non-communicable diseases as specific women’s health problems in addition to reproductive health, does suggest it could alleviate the limitations of conflating of women’s health to reproductive function. In addition, the agenda actively discourages the systematic exclusion of women from clinical research as well as advocating for interdisciplinary perspectives to be adopted in research questions regarding women’s health (Peters et al., 2016). However, the framework does not accept an integrative perspective of women’s health. It calls for the diversion of resources and attention in the global agenda for women’s health away from women’s reproductive health towards the prevention, management and treatment of non-communicable diseases in women. Justification for focusing on non-communicable diseases in women relies on both their increasing impact on morbidity and mortality accompanied by the perceived unimportance of reproductive health to women’s health and wellbeing following success in the Millennium Development Goals. The belief that reproductive health is ‘no longer as important’ does not establish an agenda for women’s health that promotes integration of reproductive health with other aspects of women’s health. Thus, the compartmentalisation of women’s reproductive function is still not addressed by the Women’s Health Agenda.

In view of the conceptual limitations in women’s health and how it is approached by public health institutes a new framework must be constructed to address such problems. The current frameworks discussed do provide some improvements to the current biomedical model. However, as shown, they cannot be considered complete solutions to the conceptual limitations of women’s health.

## 3. What can an evolutionary perspective add to the current framework of women’s health in a public health context?

### Synopsis

In this chapter I will argue a further limitation of this model of health and disease lies in its recognition of only proximate causes of disease - the biochemical and physiological mechanisms mediating health and disease during the lifetime of the individual. For a complete biological framework, this proximate perspective does not offer a complete biological explanation for health and disease (Nesse, Bergstrom et al. 2010). A complete biological explanation can only be achieved by assessing the proximate and ultimate origins of a trait - what is the trait; how did the trait develop; whether and how has it given a selective advantage; and what is its phylogeny. Thus, applying evolutionary perspectives to health will alleviate conceptual limitations in public health, especially those related to women’s health. Evolutionary perspectives on women’s health will be extended to the example of breast cancer to highlight the current conceptual limitations in women’s health and how it produces barriers to associating reproductive cancers with non-reproductive functions. From this, the association between thyroid cancer and reproductive functions can be justified.

### 2.1 Evolutionary approaches to public health

Evolutionary medicine has recently emerged as a new medical discipline within academic and research institutes. Compared to a biomedical framework of health, an evolutionary approach to health recognises patterns of frequency in phenotypes affecting health and disease within and between populations. Variation is approached from the perspective of medically significant genetic variation which may have originated from several evolutionary processes (Stearns 2012). Evolutionary processes such as random mutation, drift, migration and bottlenecks can all affect population level patterns in phenotypes through random population phenomena (Stearns 2012, Nesse, Bergstrom et al. 2010). Statistically significant changes to population phenotypes can occur through natural selection, an evolutionary process which involves the concept of reproductive success driving adaptive phenotypes. The more a trait increases the fitness of the individual, the more likely it is to be fixed in the gene pool of a population. Phenotypic traits of health that allow an individual to survive to reproductive age and successfully reproduce will be positively selected for. Health from an evolutionary perspective is therefore a means to an end: reproductive success. Medically significant genetic variation can cause differences in disease response in different populations. Health from an evolutionary perspective does not confer perfection as found in the biomedical soma. A state of non-health reduces reproductive success rather than cause manifestations of disease.

#### 2.1.1 Evolutionary approaches to public health: Life history theory

A key concept in evolutionary perspectives of health is that of the life history. Life history theory is a central tenet of evolutionary approaches to health and describes patterns of growth, development, reproduction and mortality in the individual’s life (Gluckman, Beedle et al. 2009, Ellison 2003). Additionally, if survival is assured, finite resources of energy and nutrients are available for reproduction, growth and maintenance (Abrams, Miller 2011). As reproductive success is central to adaptations in the body this means that, in face of limited resources, a trade-off between reproductive success of the individual and growth and maintenance of the body is expected. Energy investment in these three central functions of the body is not only interconnected but also reproductive success will be prioritised over growth and maintenance in adverse circumstances. The coordinated response of diverting energy to reproductive function is facilitated by hormones from the hypothalamic-pituitary-gonadal axis and thus is a physiological response (Roney 2016). Given that perfection of physiological functions is a key concept of biomedical health, a further point of discordance arises between biomedical and evolutionary health perspectives (Canguilhem, Fawcett et al. 1978). In situations of limited resources, reproductive success makes growth and maintenance of the body imperfect. Physiological processes selected for in the body can and will compromise health of the individual in favour of reproductive success in adverse circumstances. Deviation from perfect function is a physiological response to limited energy resources and does not always lead to disease as expected in the biomedical model of health.

#### 2.1.2 Application of evolutionary approaches to public health research

Given that health is determined by reproductive success and reproductive strategies differ between the sexes, it must be recognised that the state of health is inherently different between the woman and the man. Women’s health conceptualised as a deviation from the male soma as found in biomedicine is not supported by this. The man and woman’s body are subject to different selection pressures coming from different reproductive strategies dictating different trade-offs. The energetically costly characteristics of women’s reproduction - menarche, menstruation, pregnancy, parturition, lactation and menopause - result in different trade-offs compared to the different energetic costs of male reproductive strategy. The energetic allocations a reproducing woman must make are inherently different in nature and magnitude to those of a man and thus growth and maintenance of the women’s body will be subject to fundamentally different energetic limitations (Abrams, Miller 2011). The overall bodily function of the woman should be considered a separate entity to the bodily function of the man.

From a public health viewpoint, these differences in the conceptualisation of health should allow research to move beyond the tenets of biomedical perspective, namely atomism, causation and the soma, to facilitate different perspectives on population level variation. By including a life history approach to the function of reproduction, growth and maintenance, the exhaustive paradigm of biomedical ‘cause and effect’ research can be revitalised. Compartmentalisation of bodily functions and reproductive health cannot be justified given their inherent connection from an ultimate viewpoint, allowing new questions and perspectives to be approached by existing biomedical institutes.

### 2.2 Introduction to evolutionary perspectives on women’s health

To fully discuss the insights of an evolutionary paradigm in women’s health, with a specific focus on reproductive cancers, the example of breast cancer will be used. This will be used to reinforce the limitations conferred from compartmentalisation of reproductive and other physiological processes. Evolutionary models can be applied to reproductive cancers to reinforce the point that not all pathologies of the reproductive tract are inherently reproductive in nature. More importantly, the possibility of non-reproductive cancers related to reproductive function will be explored. The application of evolutionary perspective to reproductive cancers illustrates the limiting effect compartmentalisation of reproductive health has on exploring causative effects of reproductive and non-reproductive pathologies. When referring to reproductive cancer, I will use the term to refer to malignancies of the reproductive tract and related organs (e.g. breasts).

Breast cancer is the leading cause of cancer mortality in women with one in eight Western women developing a breast neoplasm during their lifetime (Strassmann 1999). Fewer than 2% of cases result from heritable mutations and thus 98% of cases are caused by environmental or life history factors (Strassmann 1999). Breast cancer and other cancers of the reproductive organs, like endometrial and ovarian cancers, have much higher incidence rates in affluent Western population than elsewhere around the world (Pollard 2008a). This incidence rate is due to mismatch between the number of menstrual cycles a contemporary Western woman will experience in comparison to women throughout hominin evolutionary history (Eaton, Pike et al. 1994, Pollard 2008b). There are two subsets of breast cancer broadly identified in women - oestrogen receptor positive (ER+) and oestrogen receptor negative (ER-). Oestrogen receptors present on a breast neoplasm indicates the tumour is sensitive to oestrogen and thus susceptible to cyclical oestrogen exposure over menstrual cycles (Aktipis, Ellis et al. 2014). The absence of oestrogen receptors as found on ER-neoplasms indicates that the neoplasm is not receptive to oestrogen levels.

#### 2.2.1 Mismatch theory

During the menstrual cycle, there are fluctuations in levels of the endogenous gonadal hormones within women - oestrogens and progesterone. Oestrogens are responsible for the proliferation of the endometrium in the follicular stage of the menstrual cycle. This elicits an inflammatory response in the endometrium, allowing the tissue to grow in anticipation of blastocyst implantation. Oestrogens are responsible for all metabolic activity in tissues involved in reproduction. Cyclical exposure of oestrogen throughout menstrual cycles will also cause an inflammatory response in tissues outside the reproductive tract. Mismatch therefore arises from the idea that the cumulative effect of oestrogen exposure throughout menstrual cycles in some contemporary western women is much greater than women throughout hominin evolution would have experienced.

**Table 3.1:** Reproductive factors affecting cumulative exposure of women to oestrogen

| Factors increasing cumulative oestrogen exposure | Factors decreasing cumulative oestrogen exposure |
| --- | --- |
| Early age of menarche | Late age of menarche |
| Later age at first birth | Early age of first birth |
| Nulliparity/Low parity | High parity |
| Use of combined hormonal/non-hormonal contraception | Use of progesterone based contraception |
| Short or no period of breastfeeding | Long periods of breastfeeding |
| Later menopause | Early menopause |

It has been postulated that the traditional number of menstrual cycles a preindustrial woman would have experienced in their lifetime is 90-130, as their fertility patterns follow those which decrease cumulative oestrogen exposure (Strassmann 1997, Eaton, Pike et al. 1994). In comparison, the number of expected menstrual cycles a contemporary western will experience in her lifetime to approach 400 (Strassmann 1997). During the menstrual cycle, ovarian hormones also stimulate cell proliferation in the breast epithelium. An increase in cell proliferation confers an enhanced risk of random genetic errors occurring in cell reproduction which may lead to carcinogenesis (Strassmann 1999). The high cumulative level of ovarian hormone exposure is thus a risk factor in the development of breast cancer in contemporary western women. Individual risk factors for breast cancer development recognised by epidemiologists include early age of menarche, nulliparity or low fertility, later age of first birth and later menopause (Aktipis, Ellis et al. 2014). The theory of mismatch arises as the environment in which adaptation occurred in hominin evolutionary history does not match with conditions found in contemporary Western societies. The high number of ovulations experienced by contemporary western women represents a departure from the reproductive pattern of most of hominin evolution (Eaton, Pike et al. 1994).

#### 2.2.2 Dysregulation of cell maintenance

The risk factors for ER-breast cancer are better understood by the evolutionary model of life history. This is a novel approach to cancer and thus there is little literature exploring this model currently. I reference a meta-analysis on social risk factors affecting ER-breast cancer which supports a life history approach. ER-breast cancer is more common amongst women of lower socio-economic status and ethnic minorities within US population (Hidaka, Boddy 2016). This suggests there is a correlation between ER-breast cancer and adverse life conditions. This supports life history theory which proposes a relationship between adverse conditions and a trade-off between reproductive function and cell maintenance. If adverse life conditions are present during early life, a ‘fast’ life history strategy may be chosen in the individual whereby energetic investment is switched from growth and maintenance to reproduction at menarche (Hidaka, Boddy 2016). Divergence of energy away from somatic maintenance can increase the risk of carcinogenesis through less investment in DNA repair and immunosurveillance in tissue beds (Casás-Selves, DeGregori 2011, Rozhok, DeGregori 2016). Additionally, survival of cells with cancerous mutations in a tissue bed is facilitated by sub-optimality of the tissue microenvironment (Rozhok, DeGregori 2016, Casás-Selves, DeGregori 2011). If tissue microenvironment is poorly maintained, there is a greater change a cancerous mutation will confer a fitness advantage over other non-mutated cells in the same tissue bed. Lack of tissue maintenance caused by divergence of energy and nutrients in those with a ‘fast’ life strategy may be a major factor in the development of ER-breast cancer in women from lower socioeconomic or ethnic minority backgrounds. This plasticity within women to adjust their reproductive strategy to an environment predicted from their early life conditions highlights the trade-off that persists within contemporary western women between reproductive function and tissue maintenance.

#### 2.2.3 Addressing conceptual limitations of women’s reproductive health

From this discussion of the risk factors of both ER+ and ER-breast cancer, I will identify a point of divergence. Through the western biomedical perspective of reproductive health - pathologies of the reproductive tract included - both ER+ and ER-breast cancers are considered reproductive cancers. However, as shown, only ER+ breast cancer has risk factors directly related to reproductive function in the woman’s lifetime. Selection for a ‘fast’ life history is hypothesised to be a risk factor for developing ER-breast cancer - this is directly related to the environment which the woman experienced in her early life. However, it is not directly related to reproductive function during her lifetime. ER+ breast cancer can be successfully treated with aromatase inhibitors, a treatment which inhibits the production of oestrogens in the body and thus reduce the proliferative properties of oestrogens in the breast neoplasm (Aktipis, Ellis et al. 2014). This treatment interferes directly with the women’s reproductive system, further promoting the association between the malignancy and reproductive function. Such a treatment would not be successful with ER-breast cancer which is more likely to be treated through chemotherapy. Thus, through this evolutionary perspective it can be argued that ER-breast cancer is not a reproductive cancer and the location of the tumour is its only association with the reproductive system.

If ER-breast cancer can be considered a non-reproductive cancer in western biomedical discourse through the application of evolutionary theory, the converse may also apply. Cancers located outside the reproductive system may have reproductive risk factors identified. As with reproductive cancers whose risk factors are all associated with the mismatch theory of modern reproductive strategies, there may be tissue beds outside the reproductive tract which respond to cyclical exposure to oestrogen in women. Through the compartmentalisation of the reproductive system in women, the association between ‘non-reproductive’ cancers and reproductive function may be overlooked or not considered when examining causative factors of the cancer. The biomedical paradigm on reproductive health may be preventing the association being made between incidence rates of ‘non-reproductive cancers’ and reproductive function in public health institutions, thus impeding preventative policies from being formed.

## 4. Applying an evolutionary framework to thyroid cancer

### Synopsis

Thyroid cancer is the most common endocrine malignancy worldwide and the fifth most common cancer in women in the USA (Nielsen, White et al. 2016, Zahid, Goldner et al. 2013, Cabanillas, Mcfadden et al. 2016). The thyroid gland, located in the neck, secretes thyroid hormones T3 and T4 which increase the metabolic rate of receptive cells (Martini, Nath et al. 2014). Control of thyroid hormone secretion is regulated by the hypothalamus-pituitary-thyroid axis through the release of thyroid stimulating hormone (TSH). Despite being classed a non-sex specific cancer, incidence rates are three times higher in women than men. I will argue that the current public health approaches to thyroid cancer do not recognise this disparity between incidence rates. Furthermore, I will show that research evaluating the impact of reproduction on women’s thyroid cancer incidence rates is limited by its conflation of parity to reproductive function. A new hypothesis relating increased risk of thyroid cancer initiation with high cumulative exposure to oestrogen will be introduced.

### 3.1 Current public health approaches to thyroid cancer

Cancer of the thyroid gland originates from the thyroid follicular epithelial cells in most cases - cells responsible for the secretion of thyroid hormones (Cabanillas, Mcfadden et al. 2016). Papillary thyroid cancer, or differentiated thyroid cancer is the most common subtype and occurs through the carcinogenesis of follicular cells. (Cabanillas, Mcfadden et al. 2016)

Thyroid cancer is of particular public health concern due to major increases in incidence rates throughout the world. In the USA there has been a 240% increase in thyroid cancer incidence rates over the past three decades (Rajoria, Suriano et al. 2012). In addition, incidence rates are three times higher amongst women compares to men with the risk of developing thyroid cancer in women being one in eight - a rate comparable to sporadic breast cancer risk (Zahid, Goldner et al. 2013, Rajoria, Suriano et al. 2010). Thus, thyroid cancer has two epidemiological points of interest: the rapid increase in overall incidence rates; and the disparity between women’s and men’s incidence rates.

From a public health perspective, differentiated thyroid cancer is associated with other thyroid disorders such as goitre, hypothyroidism and autoimmune conditions of the thyroid, such as Hashimoto’s thyroiditis (Duntas, Amino et al. 2012). Relative risk factors associated with thyroid cancer include body mass index, height, vegetable consumption, smoking, alcohol consumption, diabetes and obesity (Zhu, Zhu et al. 2016). An established causal factor of these thyroid disorders is iodine deficiency which has received considerable attention from public health authorities (Duntas, Amino et al. 2012). Iodine is a key component in T3 and T4 production with iodine deficiency decreasing the production of both hormones. Through the hypothalamus-pituitary-thyroid axis, a reduction in circulating thyroid hormones stimulates the release of TSH. TSH causes cell proliferation in thyroid follicular cells, producing chronic inflammation of the gland, characteristic of goitre.

However, the global status of iodine deficiency has improved during the 20^th^ Century with a decrease between 2003 and 2007 in the number of countries where iodine deficiency is considered a public health problem (de Benoist, McLean et al. 2008). Neither of thyroid cancer’s epidemiological phenomena can be attributed to iodine deficiency given a reduction in occurrence worldwide. Furthermore, despite continued focus on the role of iodine deficiency in thyroid disorders the WHO and American Thyroid Association there is little to no reference to thyroid cancer’s increasing prevalence and substantial differences between sex incidence rates. Another established risk factor for thyroid malignancies is ionising radiation exposure that victims of Chernobyl, Hiroshima and Nagasaki endured (Nielsen, White et al. 2016). Thyroid malignancies thus become a public health concern after nuclear disasters, but with such events incredibly rare worldwide ionising radiation cannot be considered a determinant of increasing thyroid cancer incidence rates.

### 3.2 The relationship between thyroid function and reproduction

To introduce a framework for understanding the link between thyroid cancer and reproduction, I will evaluate current evidence linking the two. By understanding the inherent association between the thyroid and women’s reproduction, I can better justify attributing thyroid cancer incidence rates to changing women’s reproductive patterns.

As mentioned, thyroid hormones regulate metabolic function throughout the body. Oestrogen increases the metabolic rate of tissues involved in reproduction such as the follicles, endometrium and breast during the follicular stage of the menstrual cycle. This similarity in function implies a direct interaction between oestrogen and the thyroid and is supported by the presence of both oestrogen receptors on thyroid follicular cells and thyroid hormone receptors on oocytes (Zahid, Goldner et al. 2013, Poppe, Velkeniers et al. 2007). This suggests the thyroid gland and gonadal axes interact continuously before and during pregnancy - thyroid disorders influence reproductive success while reproduction affects thyroid function and can influence the development of pathologies (Poppe, Velkeniers et al. 2008).

The proximate relationship between thyroid function and reproduction can be identified through the effect of recognised thyroid dysfunctions on successful reproduction. Oocyte maturation in the ovaries demands a favourable endocrine environment, including optimality in levels of T3 and T4 (Poppe, Velkeniers et al. 2007). Thus, normal thyroid function is required to produce successful oocytes and is responsible for maintaining an optimal endometrial environment anticipating implantation. The direct and indirect effect of the thyroid on normal reproduction is also evident through the impact of hypo/hyperthyroidism on success during pregnancy (Koutras 1997). Both disorders can cause menstrual disturbances in women and are both associated with reduced fertility (Koutras 1997). Indirect association occurs through thyroid mediated fluctuations in sex hormone-binding globulin (SHBG) - a glycoprotein which affects the levels of ‘free’ oestrogens circulating through the body. Increased SHBG will decrease the availability of free oestrogens available to bind to oestrogen receptors while decreased SHBG acts inversely. Hypothyroidism is associated with decreased levels of SHBG while hyperthyroidism is associated with increased levels of SHGB (Koutras 1997, Poppe, Velkeniers et al. 2008). The effect of thyroid disorders on reproduction is so prominent that successful pregnancy is rare in hypothyroidism (Koutras 1997).

The relationship between oestrogens and thyroid function can be investigated through similar means - the presence of oestrogen receptors on thyroid follicular cells and the role of oestrogen in thyroid dysfunctions. Oestrogens are steroid hormones which affect growth, differentiation and function of target cells. Given the presence of oestrogen receptors on follicular thyroid cells, proliferation mediated by oestrogens can be inferred (Rajoria, Suriano et al. 2012, Yane, Kitahori et al. 1994). The importance of ERs on thyroid neoplasms has been well documented with the recognition that neoplastic tissue contains overexpression of ERs compared to healthy tissue (Rajoria, Suriano et al. 2012). Imbalance between two subtypes of ERs, ER-α and ER-β, has also been identified on thyroid neoplasms. ER-α receptors facilitate oestrogen mediated cell proliferation and growth while ER-β receptors promote oestrogen mediated cell apoptosis and tumour suppressing functions (Rajoria, Suriano et al. 2012). Overexpression of ER-α will thus promote cell proliferation with reduced apoptosis, a characteristic of carcinogenic cell growth. The mode through which oestrogens promote cell proliferation involves the binding of a complex from the ER onto the ERE promotor region of cellular DNA, increasing transcription of growth factors (Rajoria, Suriano et al. 2012). An indirect role of oestrogen in tumorigenesis has also been implied by the inflammatory response oestrogens can cause on receptive tissue and the tumour promoting effect of chronic inflammation (Tafani, De Santis et al. 2014). Thus, oestrogens can be implicated in both the proliferation of thyroid neoplasms through direct interaction with ERs and indirectly influencing the tumour microenvironment. It must be noted, however, that many studies into the effect of oestrogen mediated tumorigenesis in the thyroid implicate oestrogens with cancer progression rather than cancer initiation.

### 3.3 Current limitations in public health research into thyroid cancer

The potential for oestrogen to influence thyroid cancer initiation and progression has not yet been recognised by health authorities such as the WHO and the American Thyroid Association. The possible link, however, has been explored in epidemiological studies. Various observational studies have been conducted to isolate specific reproductive characteristics with thyroid cancer. The studies referenced here were conducted through surveying women with thyroid cancer on several aspects of their reproductive history. Many findings have been inconsistent between studies (Table 4.1).

**Table 4.1:**
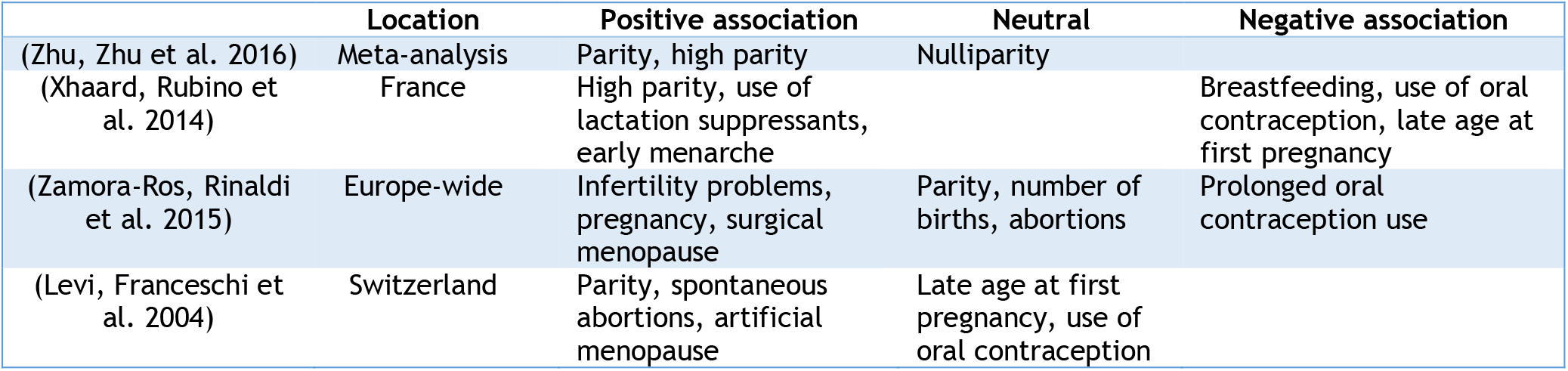
Comparison of reproductive associations found in epidemiological studies on thyroid cancer

Inferences about the link between reproduction and thyroid cancer tend to be made by linking parity to increased thyroid cancer risk. As discussed previously, equating women’s reproduction to childbirth is a simplistic, essentialised view of a woman’s reproductive function. In regards to oestrogen exposure, simply measuring parity does not explore factors such as early/late menarche, lactational amenorrheoa or use of long and short acting reversible contraception. While these metrics of reproduction have been included in some of the above studies, there is no consensus over their importance in determining the women’s experience of reproduction in these epidemiological studies observing reproductive function. For example, using oral contraceptives as a homogenous factor disregards both the use of non-oral contraceptive methods as well as equating the function of the combined oestrogen + progesterone pill and progesterone-only pill. Progesterone based contraception has a suppressive effect on oestrogen production and thus can significantly impact the oestrogen exposure in a woman. Furthermore, the associations between infertility problems, spontaneous abortions and artificial menopause and an increased risk of thyroid cancer in such studies should not be considered causative given the importance of thyroid function in successful pregnancies. Thus, infertility problems are more likely indicative of pre-existing thyroid dysfunctions progressing to thyroid cancer than infertility playing a causative role in thyroid cancer initiation.

### 3.4 Synthesising a new framework for public health research into thyroid cancer

I justify the redefinition of thyroid cancer as a women’s reproductive cancer using the evidence that thyroid cancer has a significantly higher incidence rate in women than men, and that incidence rates overall have been increasing. Mismatch theory of reproductive patterns also supports the increased incidence rates observed. As explained in the previous chapter, increased cumulative exposure to oestrogen over a contemporary western woman’s lifetime will facilitate tumorigenesis. Considering current evidence linking reproductive function to thyroid function, and the explicit link between oestrogen and thyroid tumorigenesis, applying a similar hypothesis to thyroid cancer is justified. Research into thyroid cancer causality can be approached with the hypothesis “Increased cumulative exposure to oestrogen throughout the women’s reproductive period increases the risk of developing thyroid cancer”.

This hypothesis will allow reproduction to be approached from the perspective of variations in cumulative oestrogen exposure through differences in menarche, lactation, parity and contraceptive usage. Furthermore, the association between thyroid cancer and parity can be explored through the role of increased circulating oestrogen during pregnancy interacting with the thyroid. The association of high parity with lower exposure to oestrogen does contradicts high parity as a risk factor for thyroid cancer. However, the direct interaction between pregnancy and thyroid function may prove to be a stronger determining factor of tumorigenesis than the impact of pregnancy on oestrogen exposure. This hypothesis also allows incorporation of environmental factors which affect oestrogen levels such as the role of adiposity. Obesity has been linked to increased oestrogen levels through the association between high insulin and suppressed production of SHBG in the liver (Pollard 2008b). As insulin insensitivity increases with adiposity, the resulting increase in circulating insulin decreases the production of SHBG, increasing the availability of oestrogens for binding to ERs. Given the existing association between obesity and thyroid cancer, this relationship can be studied under the hypothesis.

I have shown that in public health approaches to thyroid cancer there is no acknowledgement of the increasing incidence rates and disparity between sex incidence rates. There is sufficient available evidence to suggest an interaction between the thyroid and reproductive system, both in their involvement in optimum functioning and their role in pathologies. Limitations of current research exploring this potential link come from the equation of women’s reproduction to parity rather than including different parameters of reproduction. Subsequently, there has been no previous association between oestrogen exposure, a component of reproduction, and risk of thyroid cancer. By proposing a new hypothesis incorporating evolutionary justification for the role of oestrogen in thyroid cancer progression, I have provided a framework for these associations to be properly studied.

## 5. Conclusion

By responding to the question “What can an evolutionary perspective on women’s health provide to the current public health paradigm” I have identified the current limitations within public health and biomedical discourse. I have also demonstrated how these limitations can be alleviated through applying evolutionary theories of health. To extend this critical analysis of current public health paradigms, I have shown, using the example of thyroid cancer, how current frameworks do not allow for non-reproductive malignancies to be related to women’s reproductive function, and that even when this link is investigated there is conflation of women’s reproductive function to parity.

Within the current biomedical discourse, women’s health is essentialised to reproductive function and procreation. This essentialisation is facilitated by the emphasis biomedicine places on the body as being a functional unit of discrete, autonomous physiological entities. Reproductive essentialisation is then perpetuated in public health and political spheres due to its entwinement with demography, making population control an issue of women’s health and women’s health an issue of population control. Reproductive function is also inherently pathologised through the biomedical concept of health as a perfectly functioning soma. In biomedicine, establishing women’s reproductive function as a deviation from the norm accepts this position more than it stimulates an explanation for its pathologisation. This limited conceptual framework has been perpetuated throughout public health authorities due to the dominance of biomedicine in Western medical institutions which disallows other approaches to healthcare, such as a biocultural model, to impact public health policy formation.

With these conceptual limitations addressed, I then introduced evolutionary perspectives to health as a framework which can expand the perception of women’s health beyond reproduction and procreation. Evolutionary theory repositions health as a state of being which confers reproductive success and that optimum functioning of the body maximises reproduction rather than achieves perfection. In the context of life history theory and the different energy requirements for male and women’s reproduction, health of both sexes is affected by different energy requirements and thus should be treated as two separate entities. This allows women’s health to be reconceptualised as normal, not inherently pathological. Using the example of breast cancer, I have introduced theories of evolutionary health - mismatch theory and life history strategy - to justify the link between ER+ breast cancer incidence rates and reproductive function as well as the non-reproductive nature of ER-breast cancer. Following from these different conceptualisations of breast cancer, I introduced the possibility of non-reproductive cancer initiation being affected by reproductive function.

In the final chapter I have focused on the specific example of thyroid cancer to illustrate how a cancer outside the women’s reproductive system may be largely affected by women’s reproductive function. The current epidemiological trends of thyroid cancer - an overall increasing incidence rate and large disparities between male and women’s incidence rates - are shown to be neglected by public health authorities. The high incidence rate in women has not stimulated much response from public health authorities which can be explained by the conceptual limitations to women’s health. I have presented evidence showing the inextricable connection between the thyroid and reproductive system in women. Current epidemiological research into the link between women’s reproduction and thyroid cancer incidence has been analysed to highlight the fundamental limitation of the research - conflating women’s reproductive function to parity - which may be the cause of inconsistencies between the findings of several studies.

While I have strived for thoroughness in my dissertation, there are several limitations to this work. First, I have assumed the sex binary and its effects on health. I base the definition of woman on female primary and secondary sexual characteristics as well as the hormones associated with female reproduction. Chromosomal abnormalities or other traits of intersex individuals are not considered, as well as different genders. Secondly, there is not as wide a breadth of literature in evolutionary medicine as in other, more established, medical disciplines. Many hypotheses proposed in the field have yet to be thoroughly tested through research. While evolutionary medicine provides a new framework for conceptualising health and disease, it remains possible the hypotheses I base my arguments on may be nullified in the future.

Through this analysis, I have justified the creation of a new hypothesis which can be used to investigate the relationship between women’s reproductive function and thyroid cancer incidence rates. By summarising the link between a woman’s cumulative oestrogen exposure over her reproductive lifespan and her risk of developing thyroid cancer, I have demonstrated how women’s health must be considered as a whole, especially considering rising rates of non-communicable diseases in women worldwide. By identifying current conceptual limitations in public health approaches to women’s health, and the modes in which to remove these barriers, I have shown that exclusion of non-reproductive health matters from women’s health needs is not justifiable. Without this realisation, public health authorities will continue to approach women’s health inadequately, hindering efforts to improve health and wellbeing in all women worldwide.

1 The Soma refers to the ‘perfect’ healthy body in anthropological analysis of biomedicine

